# A Framework to enhance the Signal-to-Noise Ratio for Quantitative Fluorescence Microscopy

**DOI:** 10.1101/2025.06.03.657680

**Authors:** Suhavi Kaur, Zhe F. Tang, David R. McMillen

**Affiliations:** Department of Physical and Chemical Sciences, University of Toronto, Mississauga, Ontario, Canada

## Abstract

Single-cell fluorescence characterization has gained much attention for studying the dynamics of individual cells in human diseases such as cancer. Despite the abundance of literature on quantitative fluorescence microscopy and its advantages in measuring cell-to-cell and spatial variation over other high-throughput instruments, it lacks a concise model that one can follow to maximize the quality of images. Here, we used the signal-to-noise ratio (SNR) model to verify camera parameters and optimize microscope settings to maximize SNR for quantitative single cell fluorescence microscopy (QSFM). We determined the microscope camera’s readout noise, dark current, photon shot noise, the clock-induced charge, and validated the additive noise model for each noise source. The dark current and the clock-induced charge were both higher than reported in literature, compromising camera sensitivity. We also reduced excess background noise and improved SNR by 3-fold, by adding secondary emission and excitation filters as well as by introducing wait time in the dark before fluorescence acquisition. Additionally, our work opens new avenues for enhancing superresolution microscopy techniques such as small molecule localization microscopy (SMLM).

## Introduction

Quantitative fluorescence microscopy is a very useful technique for characterizing cells at an individual level, offering important advantages over other fluorescence measuring instruments such as the microplate reader and flow cytometer. Single cell fluorescence characterization can be used to study cellular decision-making processes that are involved in diseases such as cancer [1, 2, 3]. The microplate reader is a high-throughput fluorescence-measuring instrument with high dynamic range, speed, and sensitivity [4]. However, it is only capable of measuring the average fluorescence from all cells in any given well, so information on cell-to-cell heterogeneity is lost [5]. Studying cellular heterogeneity is important because cell-to-cell variability has even been observed in genetically identical cells [6]. The underlying cause of such variability has been attributed to stochastic noise in various cellular processes [6], such as the number of ribosomes, RNA polymerases, and gene products [7]. A flow cytometer can conduct rapid single-cell analyses for tens of thousands of cells [8]. However, this technique does not measure spatial heterogeneity within each cell. It also relies on intricate spectral unmixing algorithms in multichannel fluorescence measurements to disentangle an observed complicated emission profile into the estimated levels of each fluorophore. The need for spectral decomposition arises due to the necessary simultaneous excitation of fluorophores and simultaneous detection at multiple wavelengths. On the other hand, fluorescence microscopy measures cell-to-cell variability, spatial variability within each cell, and the level of each fluorophore in a multichannel fluorescence assay can be directly determined from the pixel intensity of the corresponding channel [9]. In this technique, multichannel fluorescence measurements do not usually involve spectral unmixing because different fluorescence channels can be determined sequentially for the same set of cells.

There is a plethora of information on different aspects of quantitative single-cell fluorescence microscopy (QSFM). For instance, a review by Jonkman et al. [9] primarily discusses sample handling factors that are important for appropriately quantifying fluorescence in cells, such as fixation, photobleaching, and sample mounting. While running QSFM experiments it is necessary to not only read biological literature but also aspects of electrical engineering. Waters et al. [10] talks about the signal-to-noise ratio and camera parameters such as noise [10], quantum efficiency, digitization, and specifics of the camera machinery [11]. However, these papers do not quantitatively analyze how different sample handling parameters or the microscope setup affect the quality of images. Knowing how those factors precisely affect the quality of images would allow us to maximize image quality given budget constraints and provide guidance on how to further improve image quality. Thus, it would be useful to place most literature on QSFM under a single theoretical framework of quantitatively optimizing signal to noise ratios and apply that framework to further improve image quality.

The standard deviation (SD) of the signal, also called total background noise (σ_total_), is contributed by the shot noise from the desired source photon (σ_photon_), the dark current (σ_dark_) [12], the clock-induced charge in an EMCCD camera (σ_CIC_) [13], and the readout noise (σ_read_) [12]. Since the different sources of noise are all independent of each other, the variance of signal (σ^2^_total_) is the sum of the variances from contributing noise sources

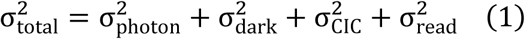

We will describe the different sources of noise, starting from the signal generating source to the recording of pixel intensity (Fig 1A). The photon shot noise refers to the noise of the incoming photons from the signal source and is also modelled by Poisson statistics [11], which describes the probability of a given number of photons striking the camera sensor within a fixed interval of time given a fixed average number of sensor-striking photons per unit time. A fraction of photons arriving at the sensor generates photoelectrons; this fraction is called quantum efficiency. In addition, heat rather than incident photons can also generate electrons that are indistinguishable from photoelectrons. The heat-generated electrons can be modelled by Poisson statistics [14] and are called the dark current. In an EMCCD camera, the electrons in the sensor, photon generated or otherwise, get shuffled through a series of cells in the gain register where entry into each subsequent cell generates additional electrons in a probabilistic manner [15]. This electron shuffling process generates additional electrons that are indistinguishable from those generated from photoelectrons [16], in a manner modelled by Poisson statistics [17]. These extra electrons are called clock-induced charge (CIC). Finally, the readout noise comes from the conversion of electrons into voltage that will eventually be converted by the Analogue-to-Digital Converter (ADC) into pixel intensity [12].

**Fig 1.**
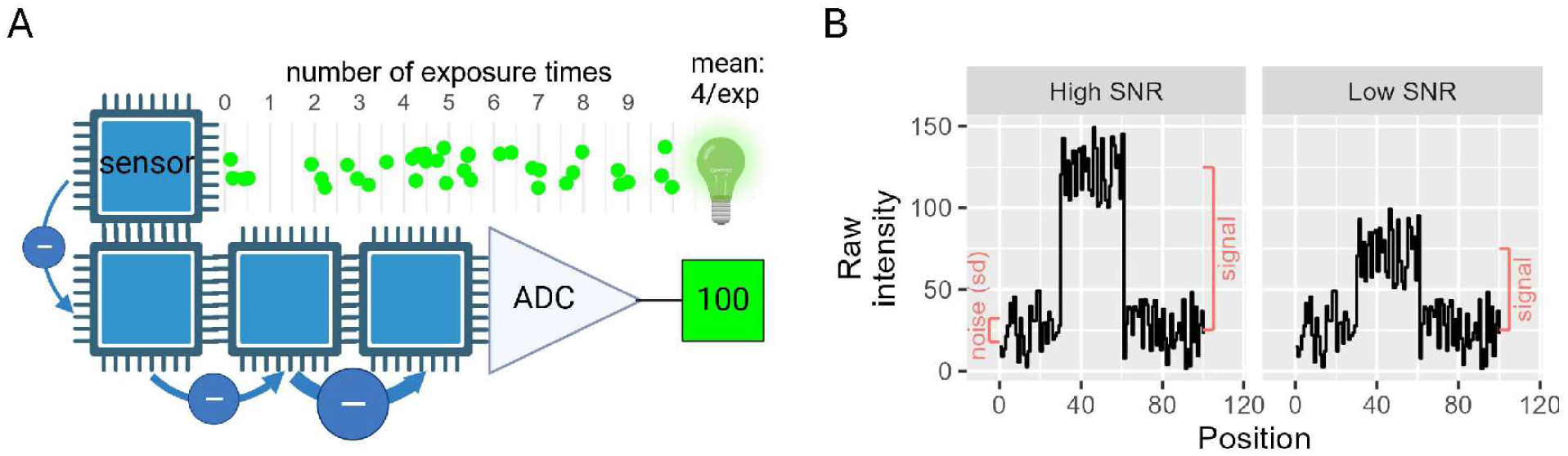
Illustrating the signal-to-noise ratio model of an EMCCD camera. (A) The path of photons (green) emitted by the light source, arriving at the camera sensor, get converted into electrons (-) that are amplified and converted into pixel intensity by the Analogue-to-Digital converter (ADC). (B) The raw intensity at high and low signal to noise ratios showing the signal of interest relative to the noise.

The electronic signal (N_e_) from the desired signal source is generated by the average number of photons (*P* × *t*) from the desired signal source that strike the camera sensor multiplied by photon to electron conversion efficiency (aka quantum efficiency QE) of the instrument [12]. Here, *P* is the average number of photons per second that comes from the signal source and *t* is the exposure time of the camera.

The SNR is the ratio of electronic signal to total noise [12]

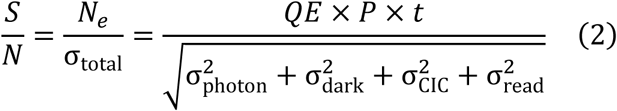

which is a measure of how much our signal of interest is above the statistical fluctuations of other signals (Fig 1B). If both signal and background emit similar intensities of light, the SNR decreases and interferes with an accurate quantification of the signal generating target.

In this study, we use the signal to noise ratio model of quantitative fluorescence microscopy to 1) verify camera parameters and 2) optimize microscope settings to maximize the signal-to-noise ratio (SNR) for QSFM. The EMCCD and sCMOS cameras are designed to reduce background and advertise low noise levels from various sources [11], costing thousands to tens of thousands of US dollars when buying them new or pre-owned. Each camera model comes with its own dark current, clock-induced-charge, and read noise specifications. However, it is difficult to tell whether the camera specifications are met from sample images of cells under bright-field or fluorescence excitation. Thus, it Is important to ensure camera parameters are within manufacturer’s specifications to maximize the value of the purchase. Additionally, expensive specialized equipment such as an EMCCD camera only results in optimal SNR if other much cheaper microscope settings are optimized to not compromise SNR. Here, we show that experimentally observed SNR could be noticeably improved to be near the theoretically maximal value permitted by the camera, by adding an extra excitation filter and an extra emission filter.

## Results

### Each camera parameter is measured by eliminating the influence of all other camera parameters

To evaluate each noise source (one of {σ_read_, σ_dark_, σ_CIC_}) and thus its corresponding camera parameter, we minimize all other noise contributing to the observed total noise *σ_total_* relative to the desired noise source. This is done to ensure that the observed total noise approximately equals the noise from the desired source. For instance, we can measure the read noise σ_read_ by taking the standard deviation of the image taken with closed light shutter to eliminate photon shot noise, 0 second exposure time to eliminate dark current noise, and no electron multiplication (EM) gain to minimize clock-induced charge. We call this image the 0 gain, 0-s exposure dark image. How other camara parameters are measured is explained in subsequent sections. It is important to note that the measured and calculated values of the camera parameters are specific to our microscope camera. However, the procedure to determine camera parameters is applicable to all camera-based microscope setups.

### Measuring pixel dependent bias and read noise

We first wanted to make sure that individual pixels do not have a systematically higher or lower value with 0 gain, 0-s exposure (0G0s) dark images. In these images, the observed noise consists only of a small amount of CIC noise at no multiplication gain and a large amount of read noise

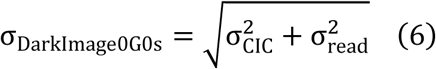

If there is no pixel dependent bias and the pixel intensity variance for different pixels are the same, then the observed variance in pixel intensity of the difference image (σ^2^_obs difference_) in a center region must equal the sum of variances of individual images (σ^2^_exp difference_)

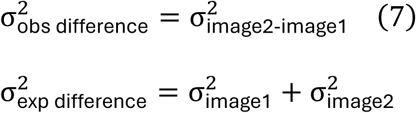

We captured five 0 gain, 0-s exposure dark images and calculated the difference images (see Methods: Data analysis) by subtracting the pixel intensity of an image at each location from the pixel intensity of the subsequent image at the same location (e.g., Image 5 – Image 4, Image 4 – Image 3, and so on). The mean expected noise within the difference image in pixel intensity (aka gray value), 42.9 ± 0.437 (mean±SD, Fig 2A) was very similar to the mean observed noise within the difference image, 41.8 ± 0.230.

**Fig 2.**
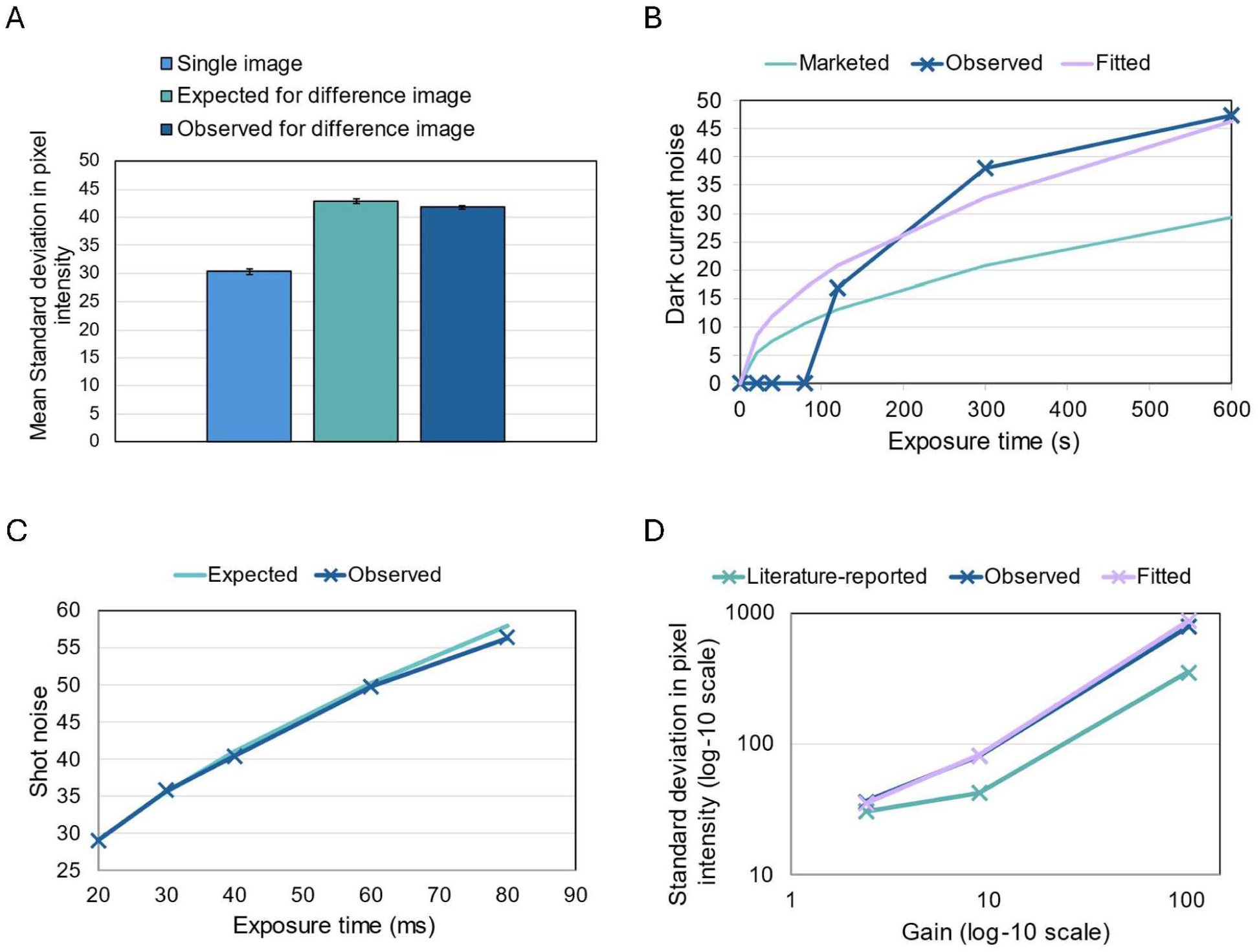
Verifying the noise model and measuring camera parameters. (A) Mean standard deviation in pixel intensity of five 0-gain, 0-s exposure dark images (light blue) and their associated difference images. (B) Dark current noise at different exposure times (0 s – 600 s). The ‘marketed’ dark current noise at different exposure times was computed based on the market dark current of 1 e-/s/pixel, using Eqn 11. The ‘observed’ dark current noise was determined from images captured with a closed camera shutter, using Eqn 10. If the dark current cannot be detected 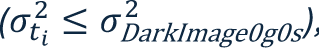, then the ti DarkImage0g0s dark current noise is set to a default value of 0. The ‘fitted’ dark current noise was computed based on the fitted dark current of 2.5 e-/s/pixel. (C) Photon shot noise of images at various exposure times (20 ms – 80 ms) with light and an open camera shutter (observed). The observed photon shot noise was calculated by removing dark image noise from the image noise using Eqn 13. The expected photon shot noise was calculated based on the observed photon shot noise at the smallest exposure time, using Eqn 14. (D) Standard deviation in pixel intensity (log-10 scale) of dark images at three different electron multiplication gains (2.40, 9.03 &103.5) on a log-10 scale (observed). The ‘literature reported’ dark image noise was calculated based on the previously reported CIC value of 4 e-, using Eqn 15. The ‘fitted’ dark image noise was calculated based on the fitted CIC value of 25 e-.

Next, we must determine the ratio between gray value (GV, aka pixel intensity) and electrons 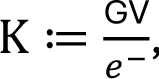, using the following relationship that is derived from the two different but equivalent ways to obtain the photon shot noise [11]

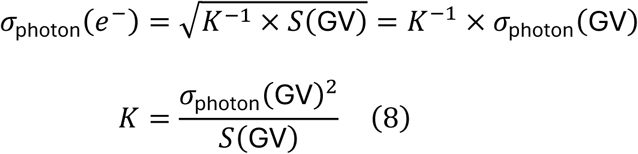

Here, signal (S(GV)) is the pixel intensity reduced by the intensity of the sample pixel in the matching dark image and σ_photon_(GV) refers to the photon shot noise in gray values that is calculated from the observed standard deviation in pixel intensity of the current image σ(GV) and of the dark image σ_DarkImage_(GV)

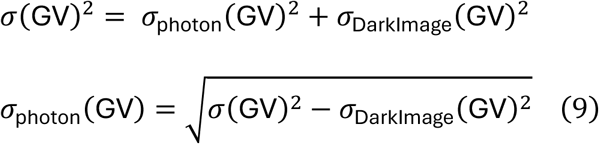

Using the gray value to electron ratio K of 1.20 and mean standard deviation in pixel intensity σ_DarkImage0G0s_ of 30.3 ± 0.577 (Fig 2A) for the 0 gain, 0-s exposure dark images, we verified that the approximate read noise of 30.3/1.20 = 25.3 *e*^-^ is within the manufacturer’s specification of 25 *e*^-^.

### Reliable extraction of the dark current

To assess the level of dark current in a camera, one may capture a series of 0 gain dark images at 0 s and multiple other exposure times (t1, t2, …) in order to isolate the dark current noise σ_*t_i_*, dark_ from the observed noise values {σ_0s_, σ_*t_i_*_|*i* ≥ 1}

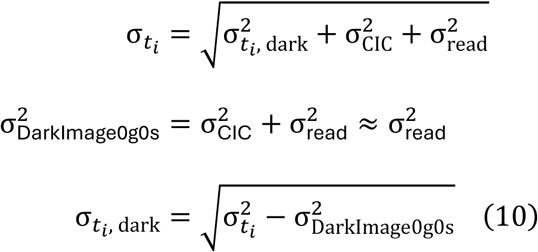

The variance of dark current is also modelled as a Poisson process, equaling to the product of the advertised dark current value (in e-/s/pixel) and the exposure time (*t* in seconds) for 0 gain dark images

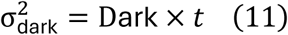

To ensure that dark noise is not a concern at short exposure times, we captured 0 gain dark images with exposure times of 20 s to 80 s and were unable to detect dark current (σ^2^_*t_i_*_ ≤ σ^2^_DarkImage0g0s_; Fig 2B). The undetectable dark current at low exposure times suggest one can spend much less on a cheaper sCMOS camera with a typical dark current of 0.1-1 e-/s/pixel [18] over an EMCCD camera with a typical dark current of <0.001 e-/s/pixel [19] and still obtain the same data quality. If a read noise of 1 e-in a sCMOS camera is satisfactory, then maximum exposure time for dark current noise to be negligible is 0.25 to 2.5 seconds

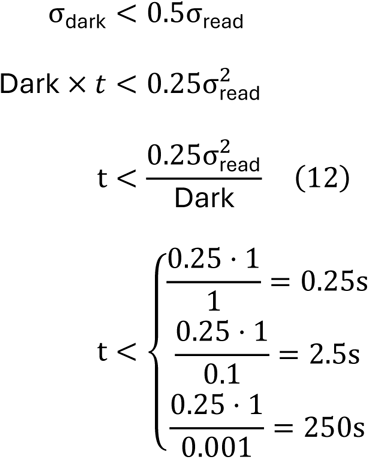

The 250 second exposure time limit for the EMCCD camera may be useful for astronomy applications, but is unnecessary for biological experiments.

Consistent with prior literature, dark current noise is revealed at high exposure times [20]. Upon capturing images at longer exposure times (120 s – 600 s) our data demonstrated a 2.5-fold higher fitted dark current of 2.5 e-/s/pixel compared to its marketed value of 1 e-/s/pixel (Fig 2B). The higher than marketed dark current value is not a serious concern, because the exposure time would need to be greater than 80 seconds for dark current noise to be noticeable. The dark current noise calculated by subtracting read noise variance had high agreement at long exposure times with theoretical noise derived from the fitted dark current value (Fig 2B), validating the additive noise model of the readout noise and dark current (Eqn 10) as well as Poisson model of the dark current noise (Eqn 11).

### Reliable extraction of photon shot noise

If two images were captured at the same setting but one image has 4 times the exposure time, then it would be expected that the isolated photon shot noise σ_4×*t*, photon_ of the image at the longer exposure time be 2 times the photon shot noise at the shorter exposure time σ_*t*, photon_. At each exposure time *t_i_*, photon shot noise σ_*t_i_*,photon_ can be isolated by subtracting dark image noise σ_DarkImage(*t_i_*_) from the observed noise σ_*t_i_*_

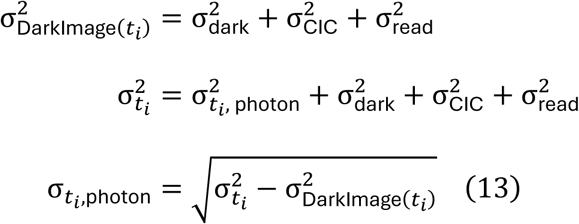

Starting from the initial exposure time *t*_1_, the photon shot noise of each subsequent exposure time *t_i_* can be calculated from the photon shot noise at that first exposure time

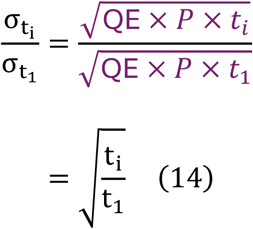

Indeed, the expected photon shot noise calculated using the above method matches the photon shot noise calculated from the observed noise at multiple exposure times (Fig 2C). This consistency validates the additive noise model of the photon shot noise σ_*t_i_*,photon_ (Eqn 13) and camera specific noise σ_DarkImage(*t_i_*)_ as well as the Poisson model of the photon shot noise (Eqn 3).

### Verifying clock-induced charge in EMCCD cameras

The clock-induced charge can be isolated from the change in observed noise between two dark images with the same short exposure time but one with no gain (*g*_O_ = 1, no EN) and the other with gain (*g_i_* > 1, EN = √2). Here, *g*_i_ refers to the gain setting in the software and is not necessarily equivalent to the EM Gain in the noise model (**Table 1**). The excess noise factor (EN) represents a statistical uncertainty introduced by the on-chip multiplication gain feature of the EMCCD camera [16].

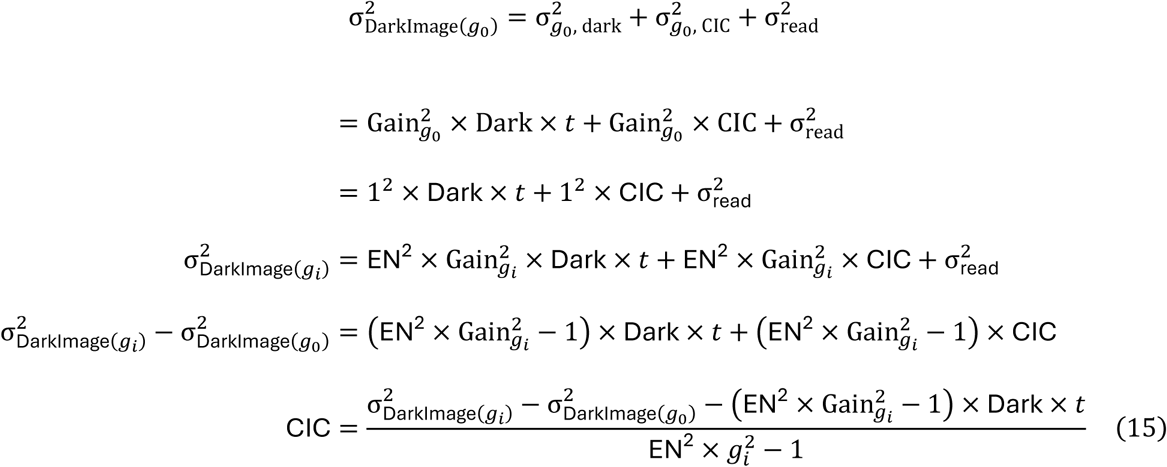

**Table 1.** The photon shot noise, dark current noise, clock-induced charge (CIC), and readout noise of an EMCCD camera for microscopy.

| NOISE SOURCE | MEAN SIGNAL | STANDARD DEVIATION |
| --- | --- | --- |
| Photon shot | $\text{Gain} \times \text{QE} \times P \times t$ | $\text{EN} \times \text{Gain} \times \sqrt{\text{QE} \times P \times t} \quad (3)$ |
| Dark current | $\text{Gain} \times \text{Dark} \times t$ | $\text{EN} \times \text{Gain} \times \sqrt{\text{Dark} \times t} \quad (4)$ |
| Clock-induced charge | $\text{Gain} \times \text{CIC}$ | $\text{EN} \times \text{Gain} \times \sqrt{\text{CIC}} \quad (5)$ |
| Readout | Arbitrary baseline | $\sigma_{\text{read}}$ |

The standard deviation formulas (black and white) are validated using our experimental data. Gain refers to the calculated electron multiplication gain, which is the ratio of the signal generated by the camera with multiplication gain compared to without multiplication gain (see Eqn 16). Dark refers to the manufacturer-reported value of dark current. The excess noise (EN) factor, quantified as 1.4 or $\sqrt{2}$ , accounts for additional variation introduced by the on-chip multiplication feature of the EMCCD camera.

We can measure the EM Gain by calculating the ratio of the signal generated by the same steady light source between the camera with multiplication gain and the camera without gain. Specifically, in addition to the two dark images taken at different gains, we also take two images of the steady light source at the same gains (*g*_O_, *g*_i_) and calculate the EM gain using the ratio of pixel intensities *I* of the four images as follows

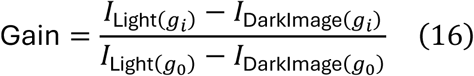

We calculated the clock-induced charge using three multiplication of gain settings and observed that average CIC is 6.25-fold higher than literature reported [17] value of 4*e*^-^. The calculated noise of the dark images using the fitted average CIC value closely matches the observed (Fig 2D), validating additive Poisson model of the CIC noise (Eqn 15)^17^. However, the expected noise of the dark images calculated using the literature reported CIC value is systematically lower (Fig 2D). At maximum EM Gain (103.5), the observed noise (σ_DarkImage(103.5)_ = 792.7) was more than 2 times greater than the expected literature-derived noise (352.3) (Fig 2D). This was an unusual observation for our Cascade 650 camera, which is marketed as having ’very high sensitivity’ and ’low noise’ under its on-chip multiplication gain feature [21], suggesting either unexpected performance degradation or a potential undetected manufacturing defect. Although dark current exceeded normal levels, it is not a concern due to the short exposure times used in biological experiments [9] (see Reliable extraction of the dark current). In contrast, the higher-than-reported CIC value cuts the signal to noise ratio by more than 50%, when high sensitivity is most needed.

### Increasing SNR by adding another emission filter

To optimize the signal-to-noise ratio (SNR), we investigated the sources of excess background noise in the acquired images of biological experiments. Excess background noise σ_*EBG*_ is any additional background noise σ_BG_ beyond observed noise of the matching dark image σ_DarkImage_

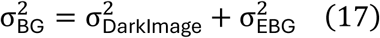

The matched dark image uses the identical parameter such as gain and exposure time but has closed shutter, such that the observed noise σ_DarkImage_ comes solely from the camera. Proper noise control would ideally make the background intensity close to dark image pixel intensity and make excess background noise relative to the dark image noise σ_*EBG*_/σ_DarkImage_ as small as possible. The excess background noise is photon shot noise according to the additive noise model of the microscope camera (Eqn 1), which is directly related to the difference in background intensity between the acquired image and the dark image according to the following

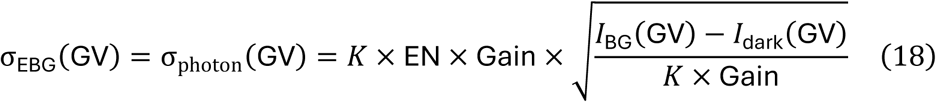

However, when measuring low-fluorescence cells on an agar pad, we observed that the relative excess background noise σ_*EBG*_/σ_DarkImage_ using the manufacturer’s recommended setup was quite high(21) (Fig 3A: OEM with cells, plot 2). In an ideal microscope setup with our noisy camera, the relative excess background noise should be a small fraction. Imaging an empty agar pad without cells resulted in 20% lower excess background noise (Fig 3A: OEM no cells, plot 2) and further removing the agar pad and the immersion oil altogether further reduced excess background noise by another 15% (Fig 3A: OEM no agar, plot 2). This suggests sample processing as a large source of excess background noise. However, the relative excess background noise is still quite high (Fig 3A: OEM no agar, plot 2), suggesting the microscope setup itself as an additional large source of excess background noise. These findings demonstrate that one should optimize SNR by prioritizing sample processing and microscope setup optimization over buying a high-end camera.

**Figure 3.**
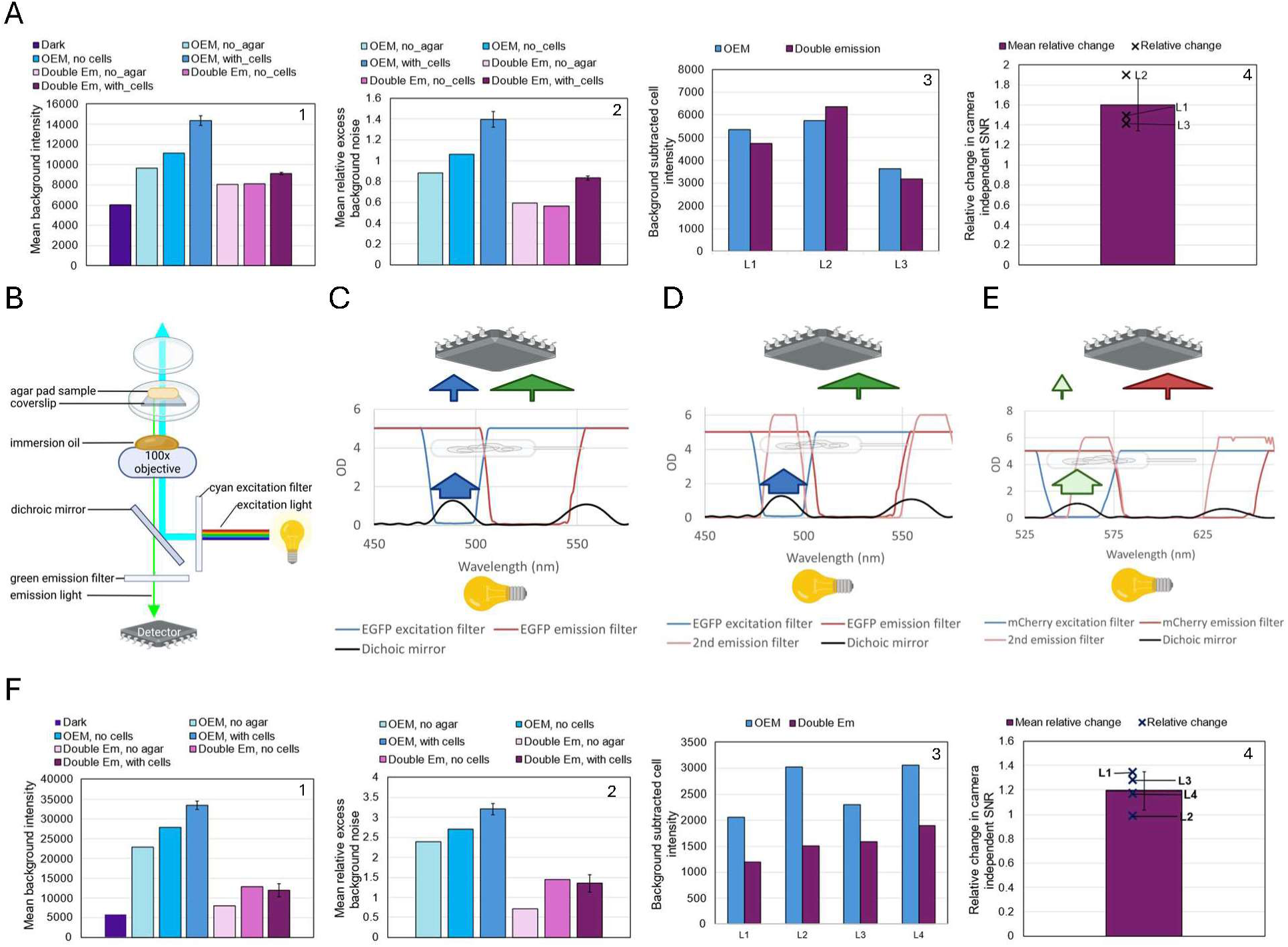
Effect of an additional emission filter on signal-to-noise ratio. (A, F) Comparing the signal to noise ratio between OEM filter setting and additional emission filter (Double Em) for EGFP (A) and mCherry (F) channels. OEM stands for the original equipment manufacturer’s microscope setup. Plots 1 and 2 show the background intensity and excess background noise relative to the dark image of images with no agar pad or oil, agar pad+oil but no cells, agar pad+oil+cells. Plots 3 and 4 show background subtracted cell intensity at 3 locations on an agar pad (4 for mCherry) and relative change in camera independent SNR of images with agar pad+oil+cells. SNR improvement is only meaningful if the improved setting shows similar or lower background subtracted cell intensity, but much lower relative excess background noise. (B) Schematic of the path of light from the fluorescence illumination system to the detector in an inverted widefield microscope using the manufacturer’s recommended setup. Created in BioRender. (C,D) Absorbance vs. wavelength spectra of EGFP single emission and excitation filters showing escaped excitation light (blue arrow) for C and blocking residual excitation light by the second emission filter for D. The width of each arrow-stem qualitatively indicates the amount of light. Created in BioRender. Note the position of the detector and light source do not reflect their true location in the microscope. (E) Absorbance vs. wavelength spectra of mCherry single excitation filter but double emission filters showing some blocking of residual excitation light but also some blocking of emission light past 625 nm.

We hypothesized that partial reflection of the excitation light by microscope optics might bypass the emission filter and reach the detector (Fig 3B), thereby increasing excess background noise. Emission filters are typically designed to transmit a specific bandwidth of light emitted from the specimen to the detector. For example, when imaging EGFP in cells, the excitation and emission filters are configured to their respective wavelengths (Ex/Em l: 488/507) [22], ensuring that only light within the desired spectral range is detected from the sample. However, the overlap in EGFP filter pair’s excitation and emission spectra at high OD [23] creates potential for leakage. Although a good emission filter is expected to block the vast majority of excitation light [24], any residual transmission of escaped excitation light contributes to background noise (Fig 3C). We introduced an additional emission filter to further block the residual excitation light from reaching the detector (Fig 3D), which enhanced the SNR by an average of 1.6-fold relative to images obtained with manufacturer’s recommended setup (Fig 3A: plot 3, 4). The SNR improvement as a result of an additional emission filter is also true for the mCherry channel, but the background subtracted cell intensity decreased (Fig 3F: plot 3, 4) due to the second emission filter only letting a much narrower set of wavelength passthrough (Fig 3E). Although we observed SNR improvement by adding another emission filter, the relative excess background noise remained high and the background intensity of the cell image and no-agar image were high relative to their matching dark image (Fig 3A: plot 2).

### Increasing SNR by increasing wait time or lowering bright-field light

We found that imaging the cells resulted in over 30% higher excess background noise than imaging without the sample or immersion oil (Fig 3A & Fig 3F), and this excess background noise was reduced by increasing waiting time in darkness or lowering intensity of the bright field light before taking the fluorescence image (Fig 4A). The >30% higher excess background noise persisted for both the OEM setup and upon addition of an extra emission filter, and is independent of the fluorescence channel (Fig 3A & Fig 3F). Increasing the wait time in the dark before fluorescence acquisition reduced the background intensity and noise at high bright-field light intensity, regardless of which fluorescence channel was used (Fig 4A). The fluorescence images taken with no agar and no immersion oil did not have bright field light immediately before fluorescence acquisition, so those images had sufficient wait time before being take and had the lowest excess background noise. However, the excess background noise was almost non-existent when the intensity of bright-field light prior to fluorescence acquisition was adjusted to a relatively low setting (Fig 4A). The reduced excess background translated to improved signal to noise ratio of fluorescent cells (Fig 4C). We wondered whether it was possible that the camera requires an unusually high amount of time to clear bright-field induced electrons before fluorescence acquisition. However, inserting a momentary darkness of 0.1 seconds between bright field and fluorescence acquisition allowed the camera to correctly register that darkness (Fig 4B). Thus, we conclude that internal partial reflections of the high bright field light within the optics system resulted in excess background noise.

**Figure 4.**
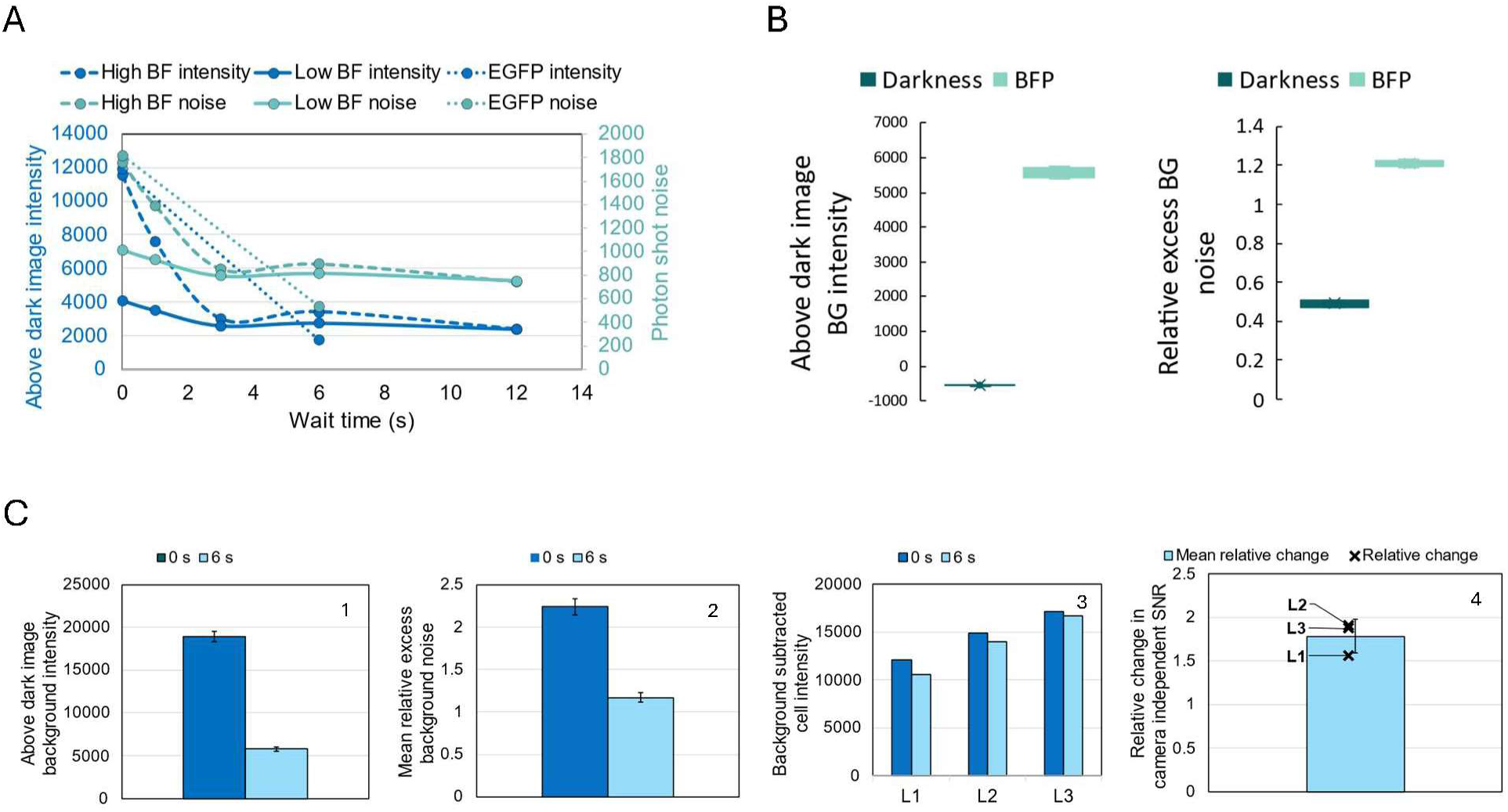
Effect of wait time or low bright field light on signal-to-noise ratio. (A) Above dark image background intensity and photon shot noise of the background for images captured at various wait times (BFP: 0 s – 12 s, EGFP: 0 s & 6 s) in darkness after bright field illumination (high and low intensity) but before fluorescence acquisition. (B) 0 s vs. 6 s wait time for BFP acquisition of E.coli cells under weak rhamnose induction of BFP. Plots 1 and 2 show above dark image back ground intensity and relative excess background noise for images taken at 3 different agar pad locations. Plots 3 and 4 show background subtracted cell intensity and relative change in camera independent SNR at 3 agar pad locations. SNR improvement is only meaningful if the improved setting shows similar or lower background subtracted cell intensity, but much lower relative excess background noise. (C) Above dark image background intensity and relative excess background noise for images taken during the transient darkness prior to fluorescence acquisition and during BFP fluorescence acquisition.

Subsequent experiments within this study automatically assumes that a suitable amount of wait time has been added before fluorescence acquisition, to minimize excess background noise induced by bright field light.

### Increase SNR by adding another excitation filter

We further narrowed down the source of excess background noise to the undesired residual emission wavelength light emitted by the fluorescence excitation source, in images that were taken with no agar and no immersion oil and double emission filters. Using our EGFP example, we hypothesized that although a good excitation filter is expected to block the vast majority of emission wavelength light, residual transmission of escaped emission wavelength light from the illumination lamp have a chance to be reflected back past the emission filter to reach the detector (Fig 5A). We introduced an additional excitation filter on top the second emission filter to remove the residual emission wavelength light from the illumination lamp (Fig 5B), and observed a further improvement in SNR (Fig 5D & Fig 5E). The combined addition of excitation filter and emission filter brought the net SNR improvement to over 3-fold.

**Figure 5.**
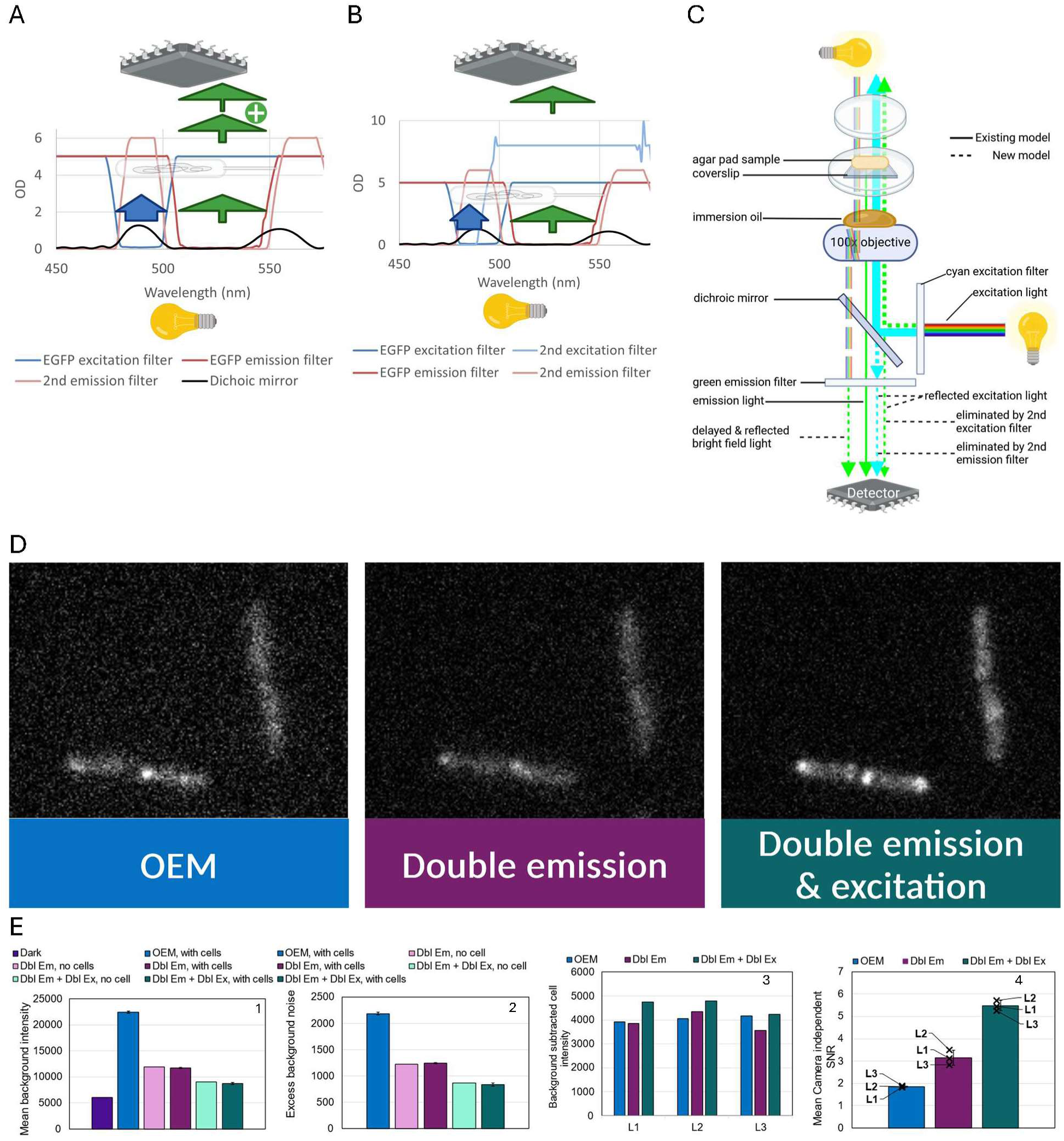
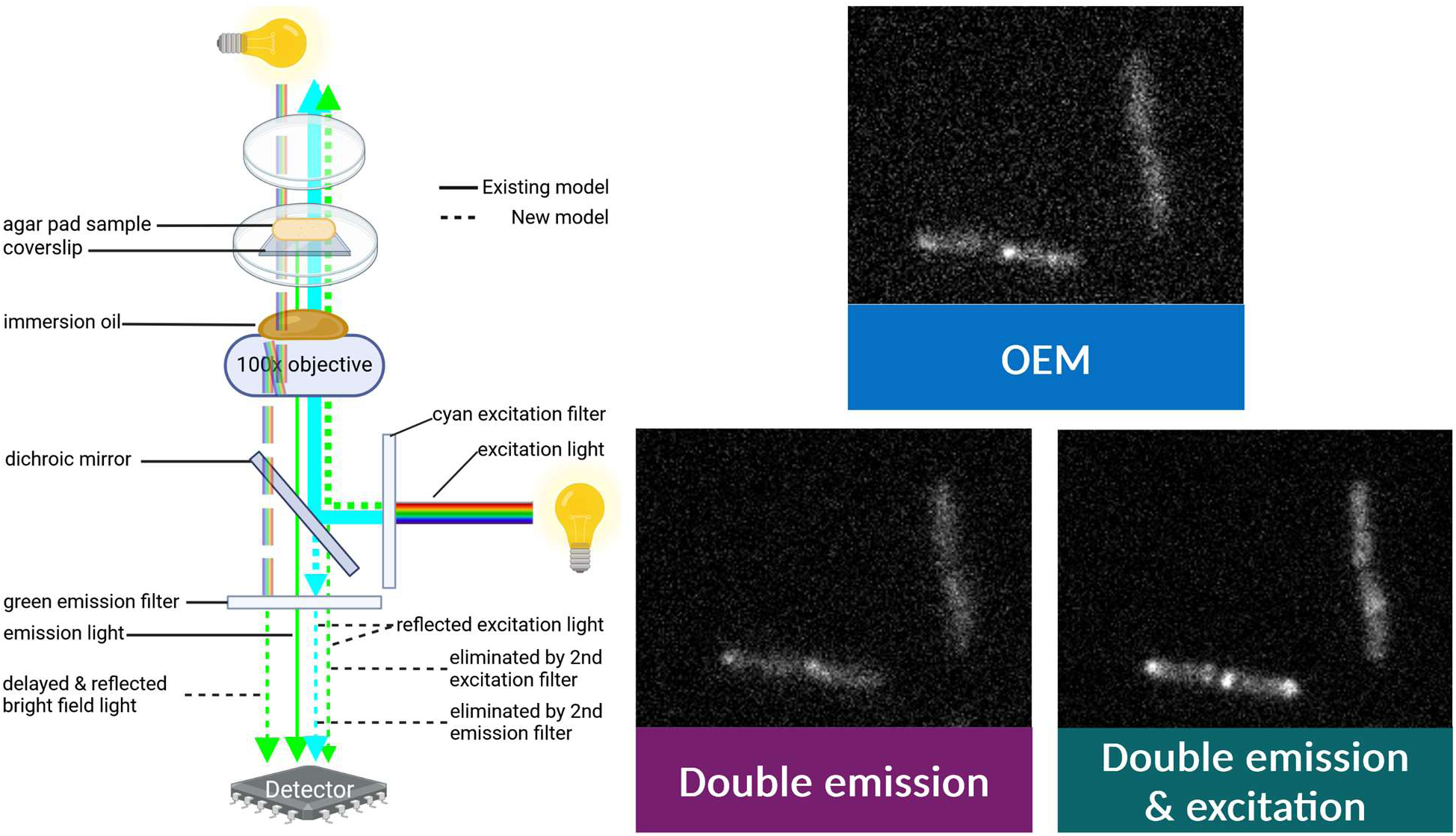
Effect of an additional excitation filter on signal to noise ratio. (A) Absorbance vs. wavelength spectra when two emission filters are blocking escaped excitation light and a single excitation filter is transmitting residual emission wavelength (green) from the illumination lamp. (B) Absorbance vs. wavelength spectra when double emission and excitation filters blocking escaped excitation light and emission wavelength from the illumination lamp, respectively. (C) The current model and new model of the path of light in a standard inverted widefield microscope setup. The new model shows escaped excitation light that reaches the detector, emission light that surpasses the single excitation filter, and reflected bright field light of the optic system. (D,E) EGFP acquisition using the OEM setup vs. double emission filters vs. double emission + double excitation filters. The images acquired under different microscope settings were processed to make them comparable: for each image, pixel intensity was subtracted by the mean pixel intensity of the background region in that image. The magnitude and amount of static in a background region reflects the background noise level. Plots 1 and 2 in panel E shows background intensity and excess background noise relative to the dark image for the dark image, agar + oil but no cell images, and agar + oil + cell images. Plots 3 and 4 in panel E shows background subtracted cell intensity and mean camera independent SNR of images at 3 different agar pad locations. SNR improvement is only meaningful if the improved setting shows similar or lower background subtracted cell intensity, but much lower relative excess background noise.

## Discussion

In this study, we analyzed the quality of microscopy images in terms of signal to noise ratio (SNR), and applied the SNR model to both verify camera parameters and optimize the microscope setup to improve SNR. We discovered that our EMCCD camera had 2.5-fold higher than marketed dark current and 6.25 fold higher than expected clock-induced charge. Although the higher dark current will not noticeably affect SNR within the typical exposure times of biological experiments, the higher clock-induced charge decreases the SNR of acquired images by more than 50% whenever high sensitivity is required the most. In addition, we discovered that SNR is lowered by high bright-field light that is partially reflected within the microscope’s optics system, by partially reflected excitation light can escape the emission filter, and by partially reflected emission wavelength light that comes from the fluorescence illumination lamp and escapes past the excitation filter (Fig 5C).

One avenue worthy of further exploration is how much SNR can be improved by our microscope setup optimization, when a newer generation of EMCCD camera is used. The lowest gain-normalized dark image noise in our EMCCD camera of

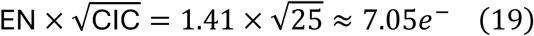

is limited by the higher than marketed clock-induced charge of 25*e*^-^. This gain-normalized dark image noise represents the true sensitivity of our EMCCD camera. On the other hand, the new generation of EMCCD cameras claim to achieve sub-electron dark image noise [19] and new generation of sCMOS cameras claim to achieve ∼1*e*^-^ dark image noise [18]. The lower noise floor matters, because it is difficult to accurately estimate the excess background noise when the excess background noise is less than 50% of the noise floor. If the excess background noise is less than 50% of the dark image noise, then the excess background variance contributes to less than 20% of total background variance

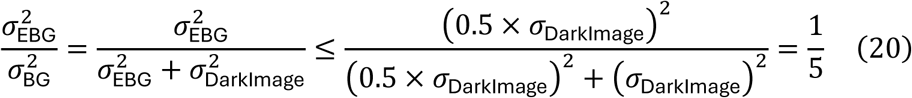

In support of our claim, we see that images with lower background than the dark image can still have ∼0.5 relative excess background noise (Fig 4B).

Our work provides a new direction to increase signal to noise ratio of microscopy images, which may enhance localization accuracy and temporal resolution of superresolution microscopy techniques. In superresolution techniques such as dSTORM, localization accuracy is strongly dependent on the intensity and accuracy of pixels within each Airy disk [25]. In MINFLUX, the temporal resolution is inversely correlated with the number of sensed electrons can be confidently attributed to a fluorophore [26]. Increasing SNR to the theoretical maximum permitted by an EMCCD camera could noticeably improve both classes of superresolution techniques.

## Materials and Methods

### Sample Preparation for image acquisition

#### Antibiotic Resistant *E. coli* Cell Culture

Three overnight cell cultures were prepared in EZ rich with either four antibiotics (4AB: 20 µg/mL chloramphenicol, 25 µg/mL carbenicillin, 15 µg/mL kanamycin, 25 µg/mL spectinomycin), or 2 antibiotic (2AB: 35 µg/mL chloramphenicol, 100 g/mL carbenicillin, 62.5 µg/mL rhamnose inducer) media. Cells were inoculated in 96-well microplates containing respective media and incubated at 37 °C with shaking.

#### Agarose Pads

A poly(dimethylsiloxane) (PDMS) chip was created using the procedure described in Ferry et al. [27] and carved with twelve (7 mm diameter) holes. The holes were used as a mold for 2% low melting point (LMP) agar pads. The PDMS chip was secured on a glass slide with tape. A solution of 4% LMP agar was synthesized with 0.20 g of LMP agar in 5 mL of distilled water, vortexed, heated in a microwave for a total of 30 s in 10 s intervals and placed in a water bath (60 °C). 1 mL solutions of 2% LMP agar were prepared with either 4AB or 2AB media in a 1:1 ratio and centrifuged for 10 s to remove air bubbles. Immediately following centrifugation, the 2% LMP agar was carefully pipetted into the PDMS molds that were placed on ice to prevent leakage of the liquid agar. A cover slip was placed over each dome to flatten the agar pads and left to solidify in the refrigerator for 30 – 45 minutes. Each mold was slightly overfilled to form a dome-shape over the top of the holes to prevent condensation between the coverslip and agar pads.

#### Agarose Pad Holder

A hole in the shape of a coverslip but with slightly smaller dimensions was carved out the lid of a petri dish. The hole was resealed with a coverslip with clear nail polish as adhesive. The solidified agar pads were inoculated with 0.25 µL diluted (1:10) cell culture and left to dry out the inoculated culture. The inoculated agar pads were placed in the petri dish with the inoculated side facing the coverslip and covered with the base of the petri dish. The final setup should resemble the sample setup in Fig 5C.

### Emission and Excitation Filter Set Up

#### OEM Setup - Single Excitation and Single Emission Filters

The inverted microscope (Nikon TE2000) was set up as per manufacturer’s instructions [28, 29] for EGFP, BFP, and mCherry acquisition via single emission and excitation filters (Chroma 86000v2).

#### Single Excitation and Double Emission Filters

An additional multi-notch emission filter (Chroma ZET405/488/561/647m) was inserted into the filter cube and the remaining setup was followed as per OEM to image EGFP, BFP, and mCherry.

#### Double Excitation and Double Emission

A second excitation filter (Chroma HQ470/40x for EGFP, DAPI EX 340-380 for BFP) was inserted in the path of the excitation light in addition to the second emission filter to image during fluorescence acquisition. Since this setup transmitted a narrower band of excitation light compared to the OEM setup, excitation light for the extra emission filter setup and the OEM setup was reduced to ensure that the transmitted excitation light achieved very similar brightness. The brightness of the transmitted excitation light that is partially reflected by the sample was measured by the camera without any emission filter.

### Image Acquisition

The petri dishes that contained both inoculated and uninoculated agar pads were subject to microscopy using 100x objective lens. No agar images were acquired without a sample or immersion oil, no cell images included oil and uninoculated agar pads, and with cells images had agar pads inoculated with respective cells. The inoculated agar pads were imaged at 3 – 4 locations. Image acquisition at each location was run by macros to limit the exposure to fluorescent light and avoid photobleaching. The order of images with different filter settings was shuffled at each location. No cell images were taken after focusing on adjacent agar pads with cells, in order to accurately measure above dark image background intensity. All microscopy images were collected using a Nikon TE2000 microscope with an attached Cascade 650 EMCCD camera. The CCD image sensor used in the Cascade 650 camera is Texas Instruments TC253.

### Data analysis

#### Image Quantification using ImageJ

A small rectangular region of interest (ROI) was selected to measure the mean pixel intensity and standard deviation of the image background. Cell intensity was measured by outlining a rough perimeter around the cell using the freehand selection tool. Images acquired from the same location but with different filter settings were stacked to maintain consistency in background and cell measurements.

Difference images were created using the image calculator feature [Process > Image Calculator > operation = Difference]. Prior to creating the difference image, a bias was introduced to one of the images [Process > Math > Add] such that there is no negative difference in pixel intensity. Any negative difference in pixel intensity is truncated to zero, making noise measurements inaccurate.

#### SNR Calculations

The photon shot noise of the background region (standard deviation due to fluorescence) was determined by subtracting the pixel intensity variance of the dark image from pixel variance of the image’s background region. Background subtracted cell intensity was calculated by subtracting the mean pixel intensity of the background region from mean pixel intensity of the selected cell perimeter. The camera-independent SNR was determined by dividing the background subtracted cell intensity by the photon shot noise of the background region.

## References

1. Albeck JG, Mills GB, Brugge JS. Frequency-modulated pulses of ERK activity transmit quantitative proliferation signals. Mol Cell. 2013 Jan 24;49(2):249–61.

2. Spencer SL, Cappell SD, Tsai FC, Overton KW, Wang CL, Meyer T. The proliferation-quiescence decision is controlled by a bifurcation in CDK2 activity at mitotic exit. Cell. 2013 Oct 10;155(2):369–83.

3. Barr AR, Cooper S, Heldt FS, Butera F, Stoy H, Mansfeld J, et al. DNA damage during S-phase mediates the proliferation-quiescence decision in the subsequent G1 via p21 expression. Nat Commun. 2017 Mar 20;8:14728.

4. Tecan Austria GmbH. Instructions for Use for INFINITE M1000 PRO. Grödig/Salzburg; 2011 Sep.

5. Bushway PJ, Mercola M, Price JH. A comparative analysis of standard microtiter plate reading versus imaging in cellular assays. Assay Drug Dev Technol. 2008 Aug;6(4):557–67.

6. Elowitz MB, Levine AJ, Siggia ED, Swain PS. Stochastic Gene Expression in a Single Cell. Science (1979). 2002 Aug 16;297(5584):1183–6.

7. Swain PS, Elowitz MB, Siggia ED. Intrinsic and extrinsic contributions to stochasticity in gene expression. Proceedings of the National Academy of Sciences. 2002 Oct 17;99(20):12795–800.

8. Nolan JP, Condello D. Spectral flow cytometry. Curr Protoc Cytom. 2013 Jan;Chapter 1:1.27.1-1.27.13.

9. Jonkman J, Brown CM, Wright GD, Anderson KI, North AJ. Tutorial: guidance for quantitative confocal microscopy. Nat Protoc. 2020 May 31;15(5):1585–611.

10. Waters JC, Wittmann T. Concepts in quantitative fluorescence microscopy. In: Methods in Cell Biology [Internet]. Elsevier; 2014 [cited 2025 Jan 4]. p. 1–18. Available from: https://www.sciencedirect.com/science/article/abs/pii/B978012420138500001X

11. Lambert TJ, Waters JC. Assessing camera performance for quantitative microscopy. In: Quantitative Imaging in Cell Biology. Boston: Academic Press; 2014. p. 35–53.

12. Mullan A. Calculating the Signal to Noise Ratio of a Camera [Internet]. 2019 Sep [cited 2024 Dec 22]. Available from: https://andor.oxinst.com/learning/view/article/ccd-signal-to-noise-ratio

13. Coates C, Mullan Alan. What is an EMCCD Camera? [Internet]. 2021 Jun [cited 2024 Dec 22]. Available from: https://andor.oxinst.com/learning/view/article/electron-multiplying-ccd-cameras

14. Merchant FA, Periasamy A. Multispectral Fluorescence Imaging. In: Microscope Image Processing [Internet]. 2nd ed. Elsevier; 2023 [cited 2024 Dec 23]. p. 201–45. Available from: https://linkinghub.elsevier.com/retrieve/pii/B9780128210499000071

15. EMCCDs: The Basics [Internet]. Teledyne Vision Solutions. 2024 [cited 2025 Jan 15]. Available from: https://www.teledynevisionsolutions.com/learn/learning-center/scientific-imaging/emccds-the-basics/

16. Fellers JT, Davidson WM. Hamamatsu Learning Center. [cited 2024 Dec 24]. Electron Multiplying Charge-Coupled Devices (EMCCDs). Available from: https://hamamatsu.magnet.fsu.edu/articles/emccds.html

17. Soesbe TC, Lewis MA, Richer E, Slavine N V., Antich PP. Development and Evaluation of an EMCCD Based Gamma Camera for Preclinical SPECT Imaging. IEEE Trans Nucl Sci. 2007 Oct;54(5):1516–24.

18. Andor Technology. Andor Technology Zyla 4.2 PLUS CL10 Water Cooled sCMOS *NEW* Microscope Camera [Internet]. 2022 [cited 2025 Feb 18]. Available from: https://www.labx.com/item/andor-technology-zyla-4-2-plus-cl10-water-cooled-scmos-new/DIS-86647-15719

19. Andor Technology. Andor Technology iXon Life 888 EMCCD *NEW* Microscope Camera [Internet]. 2022 [cited 2025 Feb 18]. Available from: https://www.labx.com/item/andor-technology-ixon-life-888-emccd-new-microscope-camera/DIS-86647-16888

20. Walters J. Sensitivity and Noise of CCD, EMCCD and sCMOS Sensors [Internet]. 2023 Apr [cited 2024 Dec 23]. Available from: https://andor.oxinst.com/learning/view/article/sensitivity-and-noise-of-ccd-emccd-and-scmos-sensors

21. Photometrics. Cascade:650 [Internet]. [cited 2024 Dec 24]. Available from: https://www.ticgroup.com.tw/menu/products/sci/ccd/cascade/650.pdf

22. FPbase. EGFP [Internet]. 2025 [cited 2025 Jan 8]. Available from: https://www.fpbase.org/protein/egfp/

23. Chroma technology. 49002 ET - EGFP (FITC/Cy2) [Internet]. [cited 2025 Jan 11]. Available from: https://www.chroma.com/products/sets/49002-et-egfp-fitc-cy2?parts=852#legend-selector1

24. Photometrics. Multichannel System Filter Selection [Internet]. 2010 [cited 2025 Jan 9]. Available from: https://www.photometrics.com/wp-content/uploads/2019/10/filters_technote.pdf

25. Lelek M, Gyparaki MT, Beliu G, Schueder F, Griffié J, Manley S, et al. Single-molecule localization microscopy. Nature Reviews Methods Primers [Internet]. 2021 Jun 3 [cited 2025 Mar 1];1(1):39. Available from: https://www.nature.com/articles/s43586-021-00038-x

26. Balzarotti F, Eilers Y, Gwosch KC, Gynnå AH, Westphal V, Stefani FD, et al. Nanometer resolution imaging and tracking of fluorescent molecules with minimal photon fluxes. Science (1979). 2017 Feb 10;355(6325):606–12.

27. Ferry MS, Razinkov IA, Hasty J. Microfluidics for synthetic biology: from design to execution. Methods Enzymol. 2011;497:295–372.

28. Nikon Inverted Microscope Instructions [Internet]. [cited 2025 Feb 5]. Available from: https://wahoo.cns.umass.edu/sites/default/files/te2000-e-u-s-m314e038cf1_1_5.pdf

29. TE2000 T-FL Epi-FL Instructions [Internet]. [cited 2025 Feb 6]. Available from: https://www.mvi-inc.com/wp-content/uploads/TE2000-EpiFl.pdf

